# Epidermal microstructures on the paired fins of marine sculpins suggest new functional hypotheses supporting benthic station-holding

**DOI:** 10.1101/2024.09.30.615682

**Authors:** Emily A. Kane, Austin M. Garner, Shubham Yadav, L. Ann Hume, Tom Pesacreta

## Abstract

Harsh environments, such as those with breaking waves and turbulent flows, present extreme challenges to organismal survival. Many animals exploiting these habitats possess adaptations to maintain position under dynamic flow conditions, such as reversible or permanent attachment systems. However, some station-holding fishes (e.g., sculpins) instead rely on morphological and behavioral modifications of their pectoral fins to increase friction with the substrate and combat drag. Despite epidermal microstructures on the fins of other benthic fishes, little exploration of pectoral fin surfaces at the microscopic scale has been undertaken in sculpins. Using scanning electron microscopy, we discovered microscopic, fibrillar projections contained within single cells on the ventral surfaces of the paired fin rays of two intertidal and two subtidal species of marine sculpins. In contrast to subtidal species, the intertidal species possessed epidermal cells with discrete channels separating groups of fibrillar projections. These features bear a striking resemblance to epidermal microstructures described in other fishes but have distinct morphological differences. We suggest the hypothesis that these previously overlooked features contribute to sculpin station-holding performance via enhanced mechanical interactions with the substrate, suggesting new taxa within which to explore potential mechanisms of underwater friction enhancement and adhesion.

## Introduction

Animals face many challenges in their environments, often possessing specialized morphological and functional adaptations to endure conditions that may otherwise limit survival. For example, aquatic environments with fast or turbulent flow can easily displace or dislodge animals and result in negative outcomes including predation, injury, and/or death. The monumental challenge of dealing with flow can thus act as an environmental filter on morphological traits, such that specific phenotypes are more common in more challenging environments [1–3]. Benthic animals subjected to strong hydrodynamic forces imposed by flow have often evolved discrete attachment systems, such as suction or adhesive-secreting discs, that help them maintain their position on aquatic substrates when exposed to flow [4–7]. However, attachment systems are not ubiquitous in turbulent environments, or even in benthic animals that maintain their position in flow. In fishes, for example, species without attachment systems maintain a stationary position (i.e., station-hold) by co-opting their body and fins to oppose or limit dislodgement forces [3,8–18]. Whole animal traits such as streamlined body shapes, small body size, and negative buoyancy are important contributors to this function, but the use of the pectoral fins, specifically, is often key to mitigating flow challenges through shape and position changes [8,13,15,17,19–21].

The pectoral fins of station-holding perciforms such as blennies (Family Blennidae), hawkfish (Family Cirrhitidae), scorpaenoid fishes such as sea robins, scorpionfishes, and rockfishes (Suborder Scorpaenoidei), and cottoid fishes including sculpins and poachers (Suborder Cottoidei) often possess a regionalized design [11,18,20,22–26]. The specialized ventral aspect of pectoral fins contacts the substrate similar to claws [18,23,24], while the more typical dorsal aspect is positioned to generate negative lift, pushing the fish toward the substrate [1,8,15]. Although the link between fin regionalization and station-holding in benthic fishes is strong, other possible mechanisms may be employed. For example, rheophilic, freshwater Ostariophysian fishes lack regionalized fins but exhibit microscopic, cornified, often elongate epithelial projections on the ventral surfaces of their paired fins, termed ‘unculi’ [27,28]. These features are characteristic of this group and function in increasing friction and/or adhesion with the substrate [29], as well as providing protection from mechanical abrasion or aid in proprioception and/or chemoreception [27,30]. Additionally, benthic darters can maintain position in flow simply by taking advantage of hydrodynamic interactions alone without the presence of regionalized fins [1,17,18,28]. Therefore, fishes likely employ a subset of possible traits to combat the forces imparted by fast or turbulent flows, and additional functional mechanisms may be present among station-holding fishes that have not been previously considered.

Despite the interest in morphological specialization for station-holding in sculpins specifically [8,11,20,23–26,31–34], existing studies primarily focus on macroscopic characteristics that aid in position maintenance. We used scanning electron microscopy to describe intra- and interspecific variation in microscopic projections on the ventral surface of pectoral and pelvic fin pads of four species of marine sculpin. For the first time, we describe raised epidermal projections similar to those described in other fishes, suggesting that such modifications may be more broadly present in benthic fishes than what has been previously described.

## Materials and methods

Three specimens from each of four species within the Superfamily Cottoidei were examined: intertidal *Artedius lateralis* and *Oligocottus maculosus*, and subtidal *Myoxycephalus polyacanthocephalus* and *Leptocottus armatus* (Figure 1, Table S1). Subtidal species represent outgroups to intertidal species, as well as the primary family Psychrolutidae, within which 3 of 4 species are contained. Phylogenetic designations follow Smith and Busby [35]. These species were chosen because they represent four different genera, show variation in contrasting streamlined or gibbose body profiles within each habitat, and individuals could be collected at similar body sizes from adjacent, shoreline habits. Although subtidal fish used in this study were juveniles, their presence within a short distance of intertidal habitats suggests they would have a higher likelihood of encountering similar flow demands as intertidal species, including the potential to get trapped in tidepools. Therefore, if there are differences compared to the adults of subtidal species, we expect that juveniles would show more exaggerated flow-related traits.

**Figure 1.**
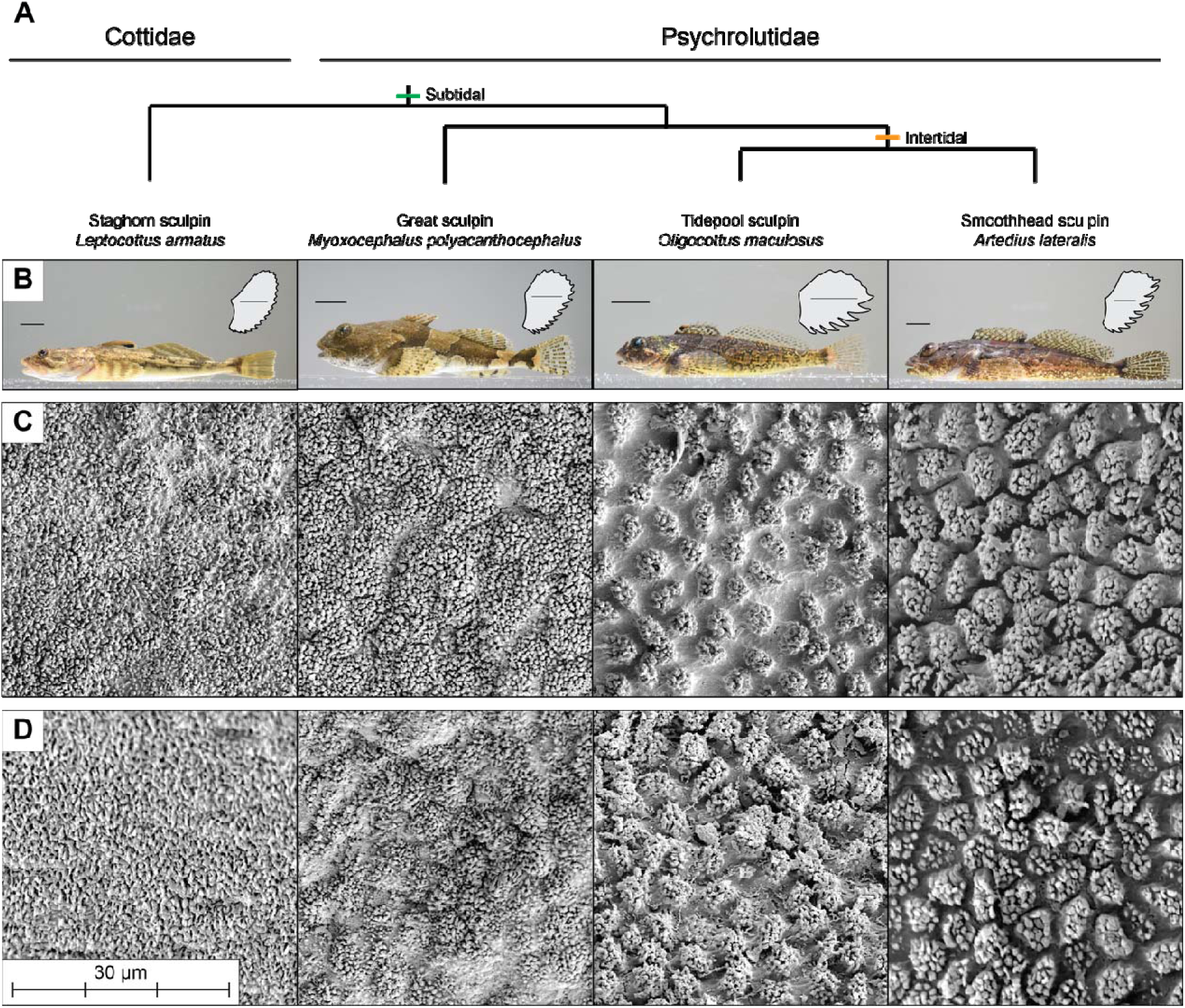
Overview of study species and results. A) Phylogenetic relationships of study species following Smith and Busby [35]. B) Lateral profile images of representative species from subtidal (green) and intertidal (orange) habitats. Images show fish collected in Summer 2022 but are not necessarily the individuals used in this work. Inset shows a tracing of a mounted pectoral fin of the individual photographed. Tracings are scaled to the same height. Scale bars are 1 cm. C-D) Representative scanning electron microscopy (SEM) images of a C) pectoral and D) pelvic fin rays of each species. Epidermal cells adorned with microscopic, fibrillar projections are present on all species. Channels separating projected areas of epidermal cells are most defined in the two intertidal species.

Individuals of each species were collected on San Juan Island, WA during the summer of 2022. Following behavioral studies not included here, fish were euthanized via immersion in tricaine methanesulfonate (MS-222), fixed in 10% formalin for 48 hours, and stored in 70% ethanol. Two specimens were selected from each species based on similarity of body size and preservation state of fins. One additional individual per species was provided by the University of Washington Burke Museum ichthyology collection (Table S1). These individuals were examined to broaden the temporal and spatial representation of samples. Museum specimens were selected to represent recent collections from sites within the Salish Sea and Puget Sound but at a distance from Friday Harbor, and at similar body sizes to the specimens collected in 2022. Unfortunately, epidermal quality of specimens obtained from the Burke Museum was poor (Figure S1), rendering data collection difficult. Data from museum specimens were excluded from further analysis, though they showed qualitative similarities to freshly collected specimens.

### Scanning Electron Microscopy

Right pectoral and pelvic fins were dissected at the base, dehydrated in 100% acetone, critical point dried, and sputter coated with a 10-12 nm gold coating. The left pectoral fin was used for specimen Alat07 because the right side was previously dissected. Samples were mounted to facilitate imaging the ventral surface of fin rays in a perpendicular plane to the electron beam. Imaging was performed with a Hitachi S3000N or Thermo Scientific Scios 2 DualBeam scanning electron microscope (SEM). Images (Hitachi: 1280 x 900 pixels; Scios 1536 x 1094 pixels) were taken at 1500X at the approximate midpoint along the free margin of each of the 5 ventral-most regionalized pectoral fin rays and the first pelvic fin ray. Some fin rays required multiple images because of the three-dimensional complexity of fin ray shape after critical point drying and the limits of tilting the stage on the SEM. Additional images were taken at lower magnifications to verify orientation and document other nearby features, but these were not used for analysis.

Following imaging, FIJI (https://fiji.sc/) was used to divide each picture into quadrants of 1000 μm each to quantify cell density and projected surface area from at least 4 quadrants per fin ray (Figure 2A). These parameters were chosen because many biomechanical (e.g., adhesion, friction) and physiological (e.g., chemo- or mechanoreception) functions are density- and area-dependent and thus may differ between species that exploit habitats with contrasting conditions. Cell density was calculated by counting the total number of cells per quadrant, including partial cells along the edges. The surface of cells contained epidermal microstructures with aspect ratios (length/width) greater than 1 and are thus hereafter referred to as fibrils. Projected surface area (hereafter referred to as projected area) of the fibrillar region of cells was determined by tracing the outline of the protruding portion (containing the fibrils) of 3 cells per quadrant. Cells for projected area measurements were selected based on the edges of the fibrillar region being visible, perpendicular to the camera view, and to encompass the shape and size variation of protrusions within the quadrant. Density and area measures were averaged across quadrants and images for each fin ray to obtain single mean values per fin ray for each individual.

**Figure 2.**
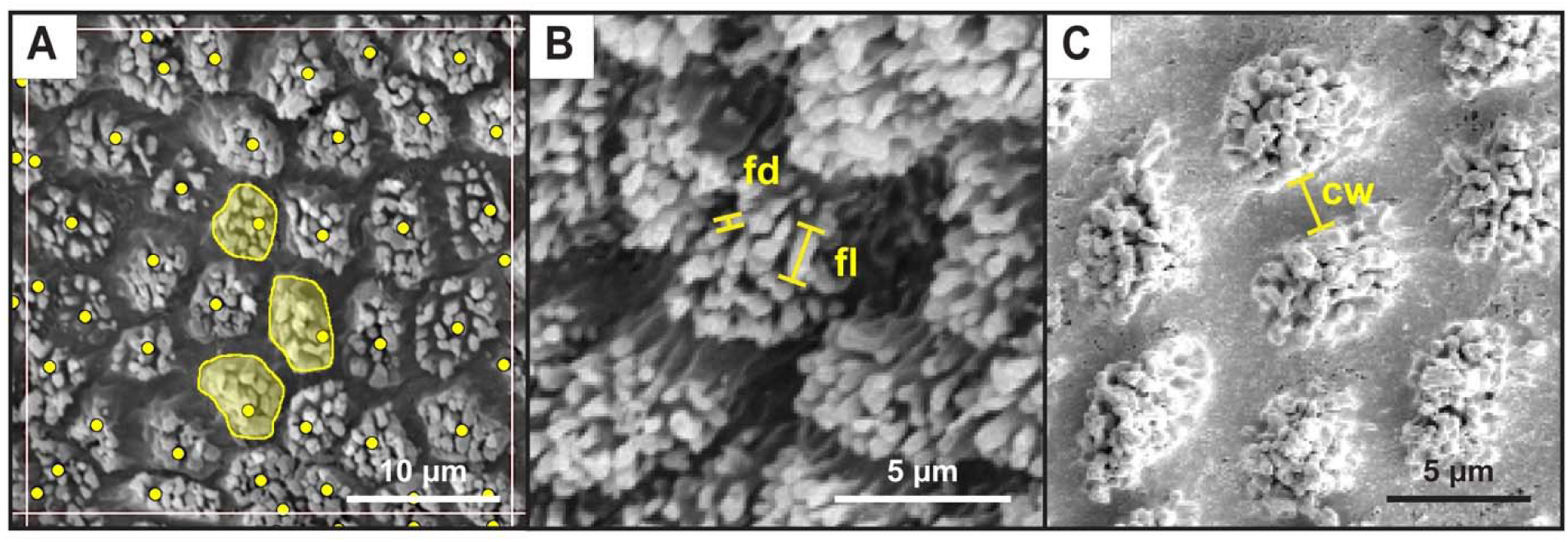
Example image analyses. A) Each grid square (bordered by white lines, only 1 shown) is 1000 μm^2^. Within each quadrant, cells were counted (yellow dots) and 3 cells/projected areas per quadrant were measured for area (yellow filled shapes). B) Linear measures of fibril length (fl) and fibril diameter (fd). C) Linear measure of channel width (cw).

To explore the morphometrics of the fibrils covering the surface of cells as well as the width of the channels between the projected area of cells, we selected the single highest quality image of the pectoral fin ray surface for each individual at 1500x magnification and made estimates of fibril density, diameter, and length, as well as channel width (Figure 2B-C). Pelvic fins were excluded for these measurements because of lower image quality. Three projected areas (or cells) per image per specimen and three fibrils per cell were selected for measurement. Fibril density was estimated by counting the number of fibrils within a specified area (usually projected area, except for *L. armatus* where cells were indistinguishable). Fibril diameter was estimated by measuring the diameter of the fibril’s tip, while fibril length was estimated by measuring the length of a line from the base to the tip of the fibril (Figure 2B). Channel width was measured as the linear distance between groups of fibers (Figure 2C). Repeat measures across cells or fibrils were averaged per individual and then an average value ± standard error calculated per species.

## Results

All four sculpin species examined displayed a ventral fin ray pad with a microtextured epidermal surface composed of fibrillar projections along the free margin of pectoral and pelvic fin rays (Figure 1). In some species, the fibrils were arranged in groups separated by distinct channels, creating a more complex, hierarchically textured surface. In contrast, the typical reticulated epidermal surface was present in areas located on the fin web, at the base of the fins, on the dorsal region of pectoral fins, and on non-primary pelvic fin rays (Figure 3). The two intertidal species (*A. lateralis* and *O. maculosus*) displayed the most prominent and easily distinguishable microtextured cells separated by channels up to approximately 2 μm in width (Table 1). Subtidal *M. polyacanthacephalus* had intermediate features that were distinguishable in some images and on some fin rays but were usually less prominent compared to the intertidal species, and channels were narrower compared to the intertidal species (Table 1). Subtidal *L. armatus* had the least well-defined features, rendering cell morphometrics (cell density, projected area, and channel width) unable to be measured.

**Figure 3.**
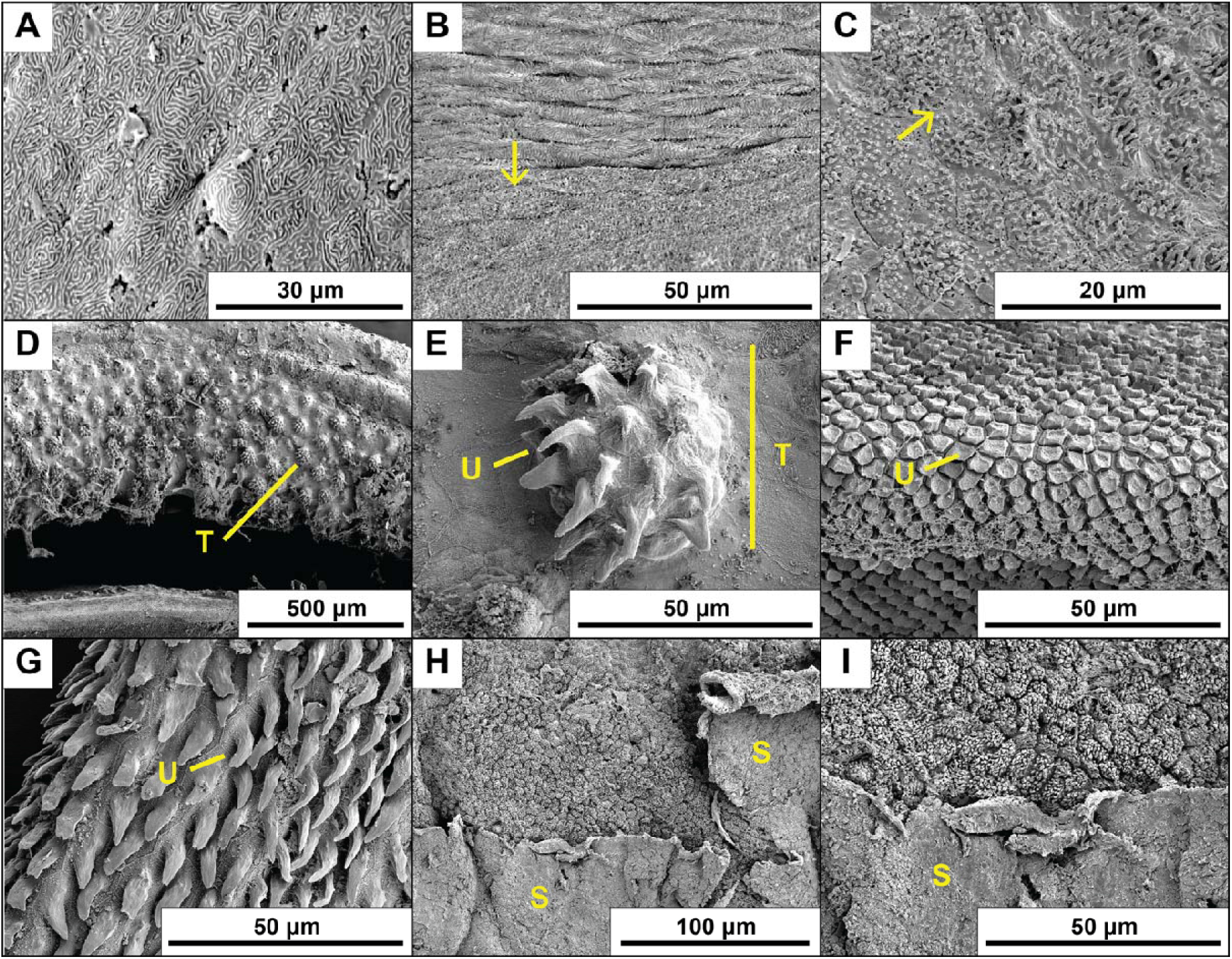
Representative images demonstrating additional observations. A) Typical epidermis from the dorsal region of the pectoral fin of *A. lateralis*. B-C) The transitional zone between typical epidermis near the pectoral fin webbing and modified epidermis on the edge of the ventral fin pad of (B) *O.maculosus* and (C) *A. lateralis.* Arrows point toward the ventral direction. D-G) Comparative images of the Ostariophysian *Paracrossochilus vittatus*: (D-E) multicellular unculiferous tubercle on the rostral cap, (F) hexagonal unculi on the inner lip, and (G) typical elongate unculi on the anterior pelvic fin ray. The specimen was a fresh preparation of a fish acquired in the aquarium trade following the methods described for sculpins. H-I) Skin sloughing observed in *M. polyacanthocephalus.* S, sloughing epidermal tissue; T, multicellular tubercule; U, unicellular unculi.

**Table 1.** Mean cell and fibril morphometrics across sculpin species examined, as well as estimates and ranges for comparative outgroups. Values from images in Figure 3 are estimates as images are not consistently perpendicular to the camera. Data are reported as means ± standard error.

| Species | Cell Density (mm<sup>-2</sup>) | Projected Area (μm<sup>2</sup>) | Fibril Density (μm<sup>-2</sup>) | Fibril Length (μm) | Fibril Diameter (μm) | Channel Width (μm) |
| --- | --- | --- | --- | --- | --- | --- |
| *Leptocottus armatus* | - | - | 1.65 ± 0.11 | 1.31 ± 0.07 | 0.58 ± 0.04 | - |
| *Myoxocephalus polyacanthocephalus* | 40,340 ± 868 | 27.01 ± 1.58 | 1.49 ± 0.12 | 1.03 ± 0.21 | 0.57 ± 0.01 | 0.71 ± 0.03 |
| *Oligocottus maculosus* | 43,121 ± 2,100 | 16.16 ± 0.74 | 1.88 ± 0.003 | 1.54 ± 0.20 | 0.49 ± 0.05 | 1.97 ± 0.16 |
| *Artedius lateralis* | 40,122 ± 2,003 | 21.52 ± 1.37 | 1.20 ± 0.03 | 2.13 ± 1.05 | 0.65 ± 0.007 | 1.59 ± 0.40 |
| Ostariophysan fishes (Figure 3) [27,29] | 5,000 – 74,000 | 7 – 29 | - | 10 – 50 | 1 – 3 | 0.17 – 8.38 |
| Clingfish [50,51,53] | 2,250 | 10,000 | - | 3 – 15 | 0.2 – 1.5 | 8 |
| Tree frogs [54,72,73] | 10,000 | 100 | - | 200 – 300 | 300 – 500 | 1 – 2 |

Overall, there was substantive variation in cell morphometrics within and across species (Table 1, Figure S2). Larger individuals generally had higher cell densities on their pectoral fins and lower densities on their pelvic fins (Figure S2). However, cell density was largely similar across species relative to the variation (Table 1, Figure S2). On average, *M. polyacanthocephalus* had the greatest projected areas, especially on pectoral fins, and *O. maculosus* had the lowest projected areas (Table 1, Figure S2). No obvious patterns were observed when comparing across pectoral fin rays relative to the variation.

Fibrils appeared to vary in density among species between approximately 1-2 fibrils per μm^2^ (Table 1). Subtidal species tended to be intermediate to intertidal species, which possessed the greatest (*O. maculosus*) and least (*A. lateralis*) dense fibrils. In contrast, *A. lateralis* had the greatest fibril length of just over 2 μm but also had the most variable fibril length compared to the other species. The smallest fibril length was just over 1 μm in *M. polyacanthocephalus*. Fibril diameter was, on average, similar across all species (Table 1).

## Discussion

Sculpins are often used as a model of station-holding morphology, behavior, and function in benthic fishes without discrete attachment devices [8,11,20,26,34,36]. We documented microscopic fibrillar projections on the ventral paired fin pads of marine sculpins that have not been previously considered or observed, despite an examination of fin ray histology in benthic fishes that included a freshwater sculpin species [28]. We hypothesize that these features are homologous to and elaborations of the typical microridges observed on epidermal tissue (Figure 3A) [reviewed in 37] based on the presence of morphological transitional states near the edges of the ventral fin pads (Figure 3B-C). Microridges are common features of vertebrate epidermal tissues, bearing anatomical and functional similarity to microvilli [37,38] but with characteristically short heights rarely more than 1 μm [38,39]. Microridges are initially formed as individual ‘pegs’ or ‘puncta’ that continuously elongate and divide laterally through fission and fusion dynamics as a result of changes in membrane tension and compression [37,40–43]. The resulting topography is highly plastic, shifting through the life of the cell [38,40,42] and responding to factors such as cell division, epidermal damage, hormones, environmental salinity, and even swimming activity level and exposure to wear [37,38,40,42–44]. Most authors suggest that their function is in mucous retention [citing 45]. Other non-mutually exclusive functions may include reinforcement of integument, protection from abrasion, increasing cell surface area, serving as an F-actin reserve, and facilitating signal transduction in response to external stimuli [37,40,43]. Given the plasticity of microridges within an individual, it seems feasible that the fibrillar projections observed on sculpin ventral fin pads may represent developmental modifications in the growth patterns of microridges. We suggest the descriptor ‘filamentous microridges’ for these features, which was used to describe a similar epidermal texture on hillstream fishes [46]. However, this terminology is contingent upon the support of structural homologies with microridges in future research.

Similar modified epidermal projections have been described on the fins and mouthparts of river-dwelling Ostariophysian fishes such as loaches and algae eaters, where they are termed ‘unculi’ [27,28]. These features are single, often hook-like, keratinous projections that extend from the surface of an epidermal cell in areas that contact the substrate (Figure 3E-G). They are exaggerated in more rheophilic species [28] and interlock with the substrate in hillstream loaches [29]. The filamentous microridges we observed in sculpins appear to be morphologically similar to the unculi on the ventral surface of the pectoral and pelvic fins of Ostariophysian fishes (Table 1). Specifically, a single, projected unculus is separated by a distinct channel or non-projected region (Figure 3F) [27]. Additionally, in some cases, unculi can have additional, smaller surface topographical features [27] similar in hierarchy to sculpin fibrils protruding from a larger, clumped structure. However, unculi on paired fins (Figure 3G) do not appear fibrillar [27,28] and are considerably wider and longer than the fibrils observed in sculpins. Further, in Ostariophysians, a single unculus adorns each cell, while multiple fibrillar projections appear to emanate from a single cell in sculpins. However, some unculi cluster together forming multicellular tubercles with discrete channels separating the clusters, as observed on the rostral cap of *Paracrossocheilus* sp. and *Garra gotyla* [27] (Figure 3D-E). These bear a resemblance to the structures we observed in sculpins except that they are multicellular clusters of unculiferous cells in these fishes and unicellular clusters of fibrillar projections in sculpins. The structures that may bear the most resemblance to sculpins are the unculi on the inner lip of *P. vittatus* (Figure 3F), which are unicellular and approximately 2 μm tall, however, they appear in a broad plateau-like morphology unlike sculpins. Reef blennies also show microstructures grossly similar to sculpins, referred to simply as a ‘cuticle’ [22,28,47,48], but it is unclear how scale and architecture compare to sculpins or Ostariophysian fishes. Unculi are often hypothesized to aid in friction enhancement. Recently, it has been demonstrated experimentally that interlocking of unculi increases conformation of ventral surfaces to rough substrates, aiding ventral suction disc performance in hillstream loaches [29]. Additionally, unculi-like structures improved pull off-force of bio-inspired suction discs compared to discs without these structures [49]. Therefore, enhanced contact with the substrate and station-holding performance is likely a significant function of these epidermal modifications in Ostariophysian fishes.

Similar to unculi, we hypothesize that the filamentous microridges observed in sculpins enhance mechanical interactions with surfaces through friction and/or adhesion, thereby contributing to station-holding performance. These fibrils and their hierarchical organization in cells with discrete channels between them are remarkably similar in their basic form to features deemed essential in generating reversible attachment in the discrete attachment systems of other aquatic organisms (Table 1). Distinct cells packed with fibrils are observed in both the adhesive suction disc of clingfish [50–52], which live in similar environments as the species studied here, and the adhesive toepads of tree and torrent frogs [reviewed in 43]. In clingfish, raised, polygonal “papillae” are divided by narrow channels that line the edge of the suction disc. Papillae are composed of bundles of hierarchical filaments that branch into nanoscopic rods. [50,51,53]. In clingfish, adhesion persists even when the suction cup seal is broken, suggesting that papillae take advantage of multiple adhesive mechanisms [51]. The adhesive toepads of tree and torrent frogs are also adorned with raised, polygonal epidermal cells separated by narrow channels and are composed of nanopillars that also have discrete channels separating them [54]. The configuration of sculpin epidermal cells appears to be most similar to that of tree and torrent frogs (with similar channel widths but smaller projected areas and greater densities) with fibrils that are more similar to the nanoscopic rods observed in clingfish (Table 1). The large differences in cellular configuration (i.e., projected area and cell density) between these groups may represent different approaches to resolving the area-density tradeoff with different morphological and/or developmental constraints rather than obvious functional differences. If cell area is constrained to small values in sculpins, then cell density must increase to cover the same epidermal surface area and vice versa in other taxa. Such observations have been made in the discrete attachment systems of other vertebrates, such as geckos and anoles, which achieve similar adhesive performance through different size and density configurations of their adhesive fibrils [55].

The epidermal cells and fibrils in sculpins likely operate using similar principles as observed or hypothesized in other taxa with discrete attachment systems. The fibrils in sculpins may contact the surface directly and interdigitate with its asperities in a process known as mechanical interlocking, similar to how the claws and setose filaments of rheophilic insects, loach unculi, clingfish papillae, and tree frog epidermal cells may interact with surfaces [6,49,56,57]. Alternatively, or additionally, projections may increase friction with the substrate through adhesive interactions. Fish epidermis is typically coated in mucus, and perhaps especially so for mostly scaleless fishes such as sculpins [58]. Thus, increases in surface area by microscopic fibrils may enhance adhesion with the substrate via capillary or viscous forces, similar to what has been suggested in clingfish papillae or tree frog toepads [51,reviewed in 54]. Channels between epidermal cells evident in intertidal sculpins may facilitate evacuation of excess mucus from the interface, decreasing the separation distance between the projections and substrate and increasing adhesive stress, similar to the mechanism observed in the papillae of clingfish suction discs [51] and unculi-like projections of a bio-inspired suction cup [49]. These hypothesized mechanisms, however, will require future biomechanical work exploring the tribological properties (adhesion, friction, wear) of sculpin fin rays.

The microstructures observed in sculpins may also serve other or non-mutually exclusive functions. Ostariophysian unculi, for example, have been implicated in chemo- or mechanoreception, feeding, mechanical protection, or antifouling [27,29,59,60]. Sensory functions are unlikely in sculpin structures as sensory receptors such as tastebuds are larger, multicellular, or with distinct morphology unlike what is observed here [28,30,61,62]. Additionally, sensory receptors and unculiferous cells occur in non-overlapping locations in Ostariophysian fishes [28], suggesting mutually exclusive functions. The presence of shorter, more robust unculi on the lips of fishes such as algae eaters (Figure 3F) suggests that mechanical abrasion may assist with feeding [27]. However, this is less likely to be an important function of fin-based epidermal microstructures, especially in suction-feeding fish such as sculpins [63–65]. Epidermal features may also provide mechanical protection from scrapes on rough substrates or from pathogens or other parasites. On some sculpin specimens, we observed sloughing of the outer epidermal layer (Figure 3H-I, S3) suggesting that the outermost layer is shed periodically. We made a similar observation more recently on the pectoral fins from freshly euthanized *O. maculosus* specimens, where these folds appeared to have a different consistency than mucous and support the conclusion that this tissue is epidermal (Figure S3). Skin sloughing has also been observed in the sculpin *Blepsias bilobus* in response to epibiotic growth [58] and in the unculiferous tissues of riverine Ostariophysian fishes, where the most well-developed unculi are present on the outermost layer following keratinization [28]. Epidermal microstructures are also reinforced in blennies [48,66], suggesting resistance to wear. In sculpins, the outer layer appears less developed and potentially worn (Figure 3H-I), suggesting that epidermal features may not be reinforced in sculpins but further work is needed to asses composition. In either case, microstructural wear likely reduces the potential of these structures to enhance surface interactions. Thus, replacement through sloughing would be beneficial for both maintaining mechanical interactions but also protecting deeper tissue. So far, we have not seen functional tests of these additional hypotheses in any fish known to have epidermal microstructures, suggesting a new area of in need of future inquiry.

We found considerable variation in the configuration and morphometrics of both the epidermal cells and fibrillar projections in marine sculpins that suggest a greater functional capacity in intertidal compared to subtidal species. Most notably, the epidermal cells bearing fibrillar projections appear to be separated by discrete channels that are widest in the environmentally more extreme intertidal species. Wider channels in intertidal fishes may facilitate mucus evacuation for enhanced contact with the substrate through mechanical or adhesive interactions relative to subtidal fishes [51], which may be more critical in the chaotic intertidal zone. The intertidal species also appear to have smaller projected areas compared to the subtidal *M. polyacanthocephalus*, likely a consequence of the wider channels but similar cell densities. Preliminarily, fibrils appear to be longer in the intertidal species compared to the subtidal species, which may increase surface area for adhesive interactions or enable enhanced conformation with surface asperities through differential bending properties [29,51]. These differences in the morphometrics of the fibrils should be considered preliminary as specimen image quality and conditions were not ideal for these measures. For example, projection densities are likely underestimated and projection diameters are likely overestimated because fibrils may be coated in residual mucus, rendering it difficult to know whether measures were of multiple fibrils clumped together or fibrils with high morphological variability. Fibril lengths are likely underestimated as it was difficult to orient the projections perfectly parallel to the electron beam. Future work will need to optimize specimen preparation or employ different imaging techniques (e.g., transmission electron microscopy) to better estimate morphometrics of these structures. Additionally, a wider range of species and individuals should be examined to further describe the potential specialization of intertidal species. Although morphological differences between species may be adaptations to the dynamic and extreme intertidal zone, invasion into intertidal habitats is a derived condition among marine sculpins that occurred once [2,35,67–69]. Thus, it is unclear whether these microscopic traits are associated with evolutionary history, environmental demands, or some combination of these factors. Nevertheless, our findings inspire new questions regarding how these features compliment other adaptations for benthic station-holding in sculpins, where traits such as small body size, gripping fin rays, and/or negative lift generation have been described as primary mechanisms [2,8,11,18,20,24,26,32,69–71].

Our observations in marine sculpins suggest a new, potentially convergent example of a morphological feature similar to structures hypothesized or demonstrated to enhance friction and/or adhesion in other aquatic vertebrates. This result represents a novel insight into the natural history of sculpins that is broadly applicable to further studies of organismal performance, attachment systems, and convergent evolution. This occurrence stretches the boundaries of our knowledge of benthic station-holding in fishes and highlights the need for inclusion of more taxa, including sculpins, in future research on friction enhancement and adhesion in biological systems.

## Acknowledgements

The authors thank Friday Harbor Lab, the Karel F. Liem Bioimaging Facility, and Kayla Hall for their assistance with preliminary work where these microscopic traits in sculpins were discovered. We also thank undergraduate Kelsey Fell who assisted with initial imaging and data analysis. Funding was provided by the University of Louisiana at Lafayette startup funds (EAK), a Friday Harbor Lab New Faculty Fellowship (EAK), and the Louisiana Board of Regents (contract number LEQSF(2023-26)-RD-A-24, EAK).

## Data accessibility

All original and analysis images and data will be available on Dryad following publication. Alcohol preserved specimens and SEM preparations will be maintained in the Kane Lab at the University of Louisiana at Lafayette.

## Ethics statement

All procedures involving the collection and housing of animals prior to preservation were approved by the Institutional Animal Care and Use Committee at the University of Washington (protocol #4238-18).

## Artificial Intelligence (AI) declaration

Artificial intelligence was not used in the preparation of this manuscript.

