## Supplemental information for "Epidermal microstructures on the paired fins of marine sculpins suggest new functional hypotheses supporting benthic station-holding"

**Table S1** Specimens used in this study. Within a species, individuals are ordered by their imaging sequence. Local samples have not been accessioned and are identified using individual IDs. Collection locations on San Juan Island include: Cattle Point (48.45422, -122.96251), Eagle Cove (48.46171, -123.0326), Deadman Bay (48.51336, -123.14686), or Jackson Beach (48.51982, -123.01101). Specific location was not tracked for each individual.

| **Species** | **Individual or lot ID** | **Collection date** | **Collection location** | **Standard length (mm)** |
| --- | --- | --- | --- | --- |
| *Leptocottus armatus* | Larm05 | 07/2022 | Cattle Point, Jackson Beach, or Eagle Cove | 86.96 |
|  | Larm04 | 07/2022 | Cattle Point, Jackson Beach, or Eagle Cove | 86.72 |
|  | UW155816 | 09/2015 | 47.95, -122.233333 | 78.55 |
| *Myoxocephalus polyacanthocephalus* | Mpol04 | 07/2022 | Cattle Point or Jackson Beach | 71.61 |
|  | Mpol02 | 07/2022 | Cattle Point or Jackson Beach | 85.10 |
|  | UW156405 | 08/2016 | 48.14139, -123.42876 | 74.57 |
| *Oligocottus maculosus* | Omac02 | 07/2022 | Cattle Point or Deadman Bay | 58.31 |
|  | Omac03 | 07/2022 | Cattle Point or Deadman Bay | 51.48 |
|  | UW156016 | 09/2015 | 47.95395, -122.284333 | 45.99 |
| *Artedius lateralis* | Alat07 | 07/2022 | Cattle Point or Deadman Bay | 78.45 |
|  | Alat02 | 07/2022 | Cattle Point or Deadman Bay | 86.35 |
|  | UW118838 | 08/2009 | 47.570405, -122.618323 | 81.11 |

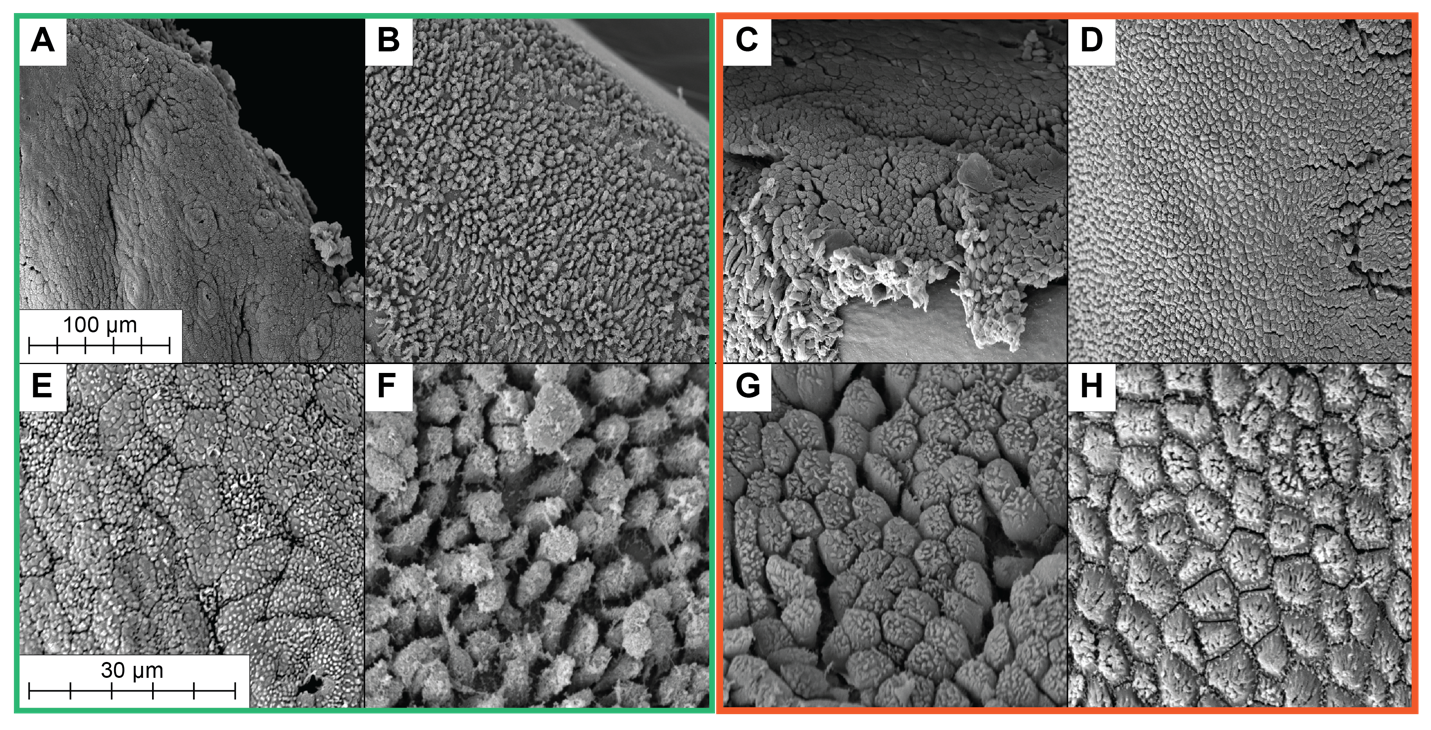

**Figure S1** Representative images of a pectoral fin ray from museum-prepped specimens of A, E) *Leptocottus armatus*, B, F) *Myoxycephalus polyacanthocephalus*, C, G) *Oligocottus maculosus*, and D, H) *Artedius lateralis*. Subtidal species are outlined in green; intertidal species are outlined in orange. A-D) Images taken at 300x showing the wider view around E-H) close-up images taken at 1500x. These images represent the best for each specimen, and many fin rays did not have any remaining epidermis. The smooth subdermis can be seen in image C.

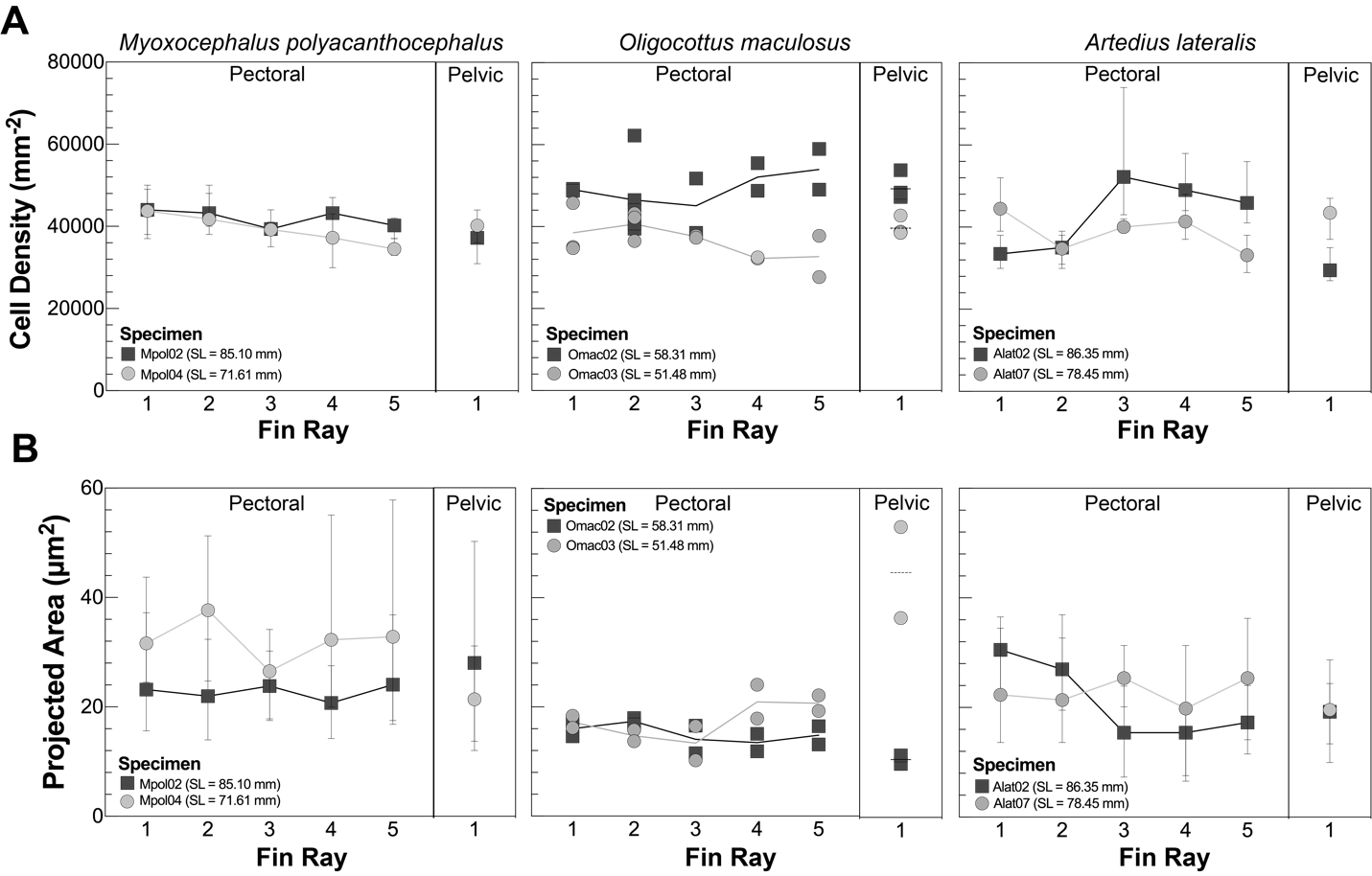

**Figure S2** A) Cell density and B) projected area measures across species, fins, and fin rays. Pectoral fin rays are ordered starting from the ventral-most ray. Density and area measures for *M. polyacanthocephalus* and *A. lateralis* are means calculated across all measured quadrants and images. Error bars represent the range of repeat measurements made per individual. For *O. maculosus*, each fin ray was imaged along multiple regions of the free margin to explore variability along fin rays. Each data point indicates the mean cell density or projected area across measured quadrants taken at different locations along each fin ray per individual. All three species show similar cell densities. Projected areas are greatest in subtidal *M. polyacanthocephalus*, lowest in *O. maculosus*, and intermediate in *A. lateralis*.

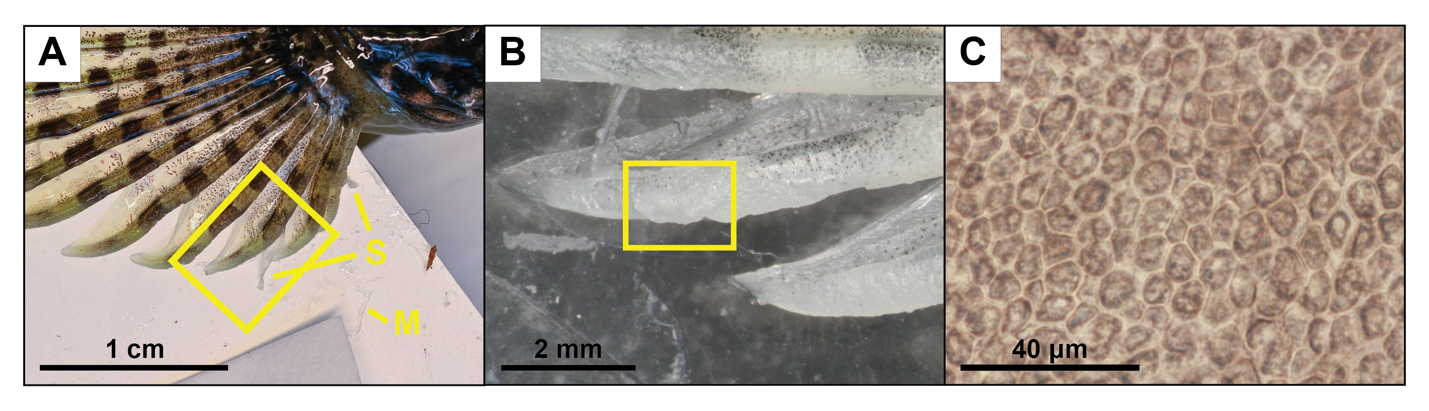

**Figure S3** Skin sloughing on an *O. maculosus* specimen. This fish was collected in July 2024 from a similar location as those used in this study and has not been used for scanning electron microscopy preparations. A) DSLR camera image of the right pectoral fin mounted for morphometric analyses. Sloughing was observed on both pectoral fins, but was more prominent on the right fin. Sloughed epidermis has a distinct color and texture compared to sloughed mucous. B) Dissecting microscope image of the formalin-preserved fin showing tissue covering the underlying epidermis. This outer covering was readily affixed to the fin ray and had to be removed by pulling. C) Stacked compound microscope image (3 images) of the removed tissue from B. Cellular structure is clearly visible, confirming that it is epidermal in origin. Cells are similar in size as those on the specimens examined here. The tissue is only 1 cell layer thick, similar to the sloughed portions shown in Figure 3. M, mucous; S, sloughing epidermal tissue.
